# First animal source metagenome assembly of *Lawsonella clevelandensis* from canine external otitis

**DOI:** 10.1101/2023.12.17.572052

**Authors:** Adrienn Gréta Tóth, Norbert Solymosi, Miklós Tenk, Zsófia Káldy, Tibor Németh

## Abstract

External otitis is one of the most common conditions in dogs to be presented to the veterinarian. Moreover, the disorder is often difficult to manage. The range and role of microorganisms involved in the pathogenesis are currently not fully understood. Therefore, the condition has been studied using third-generation sequencing (Oxford Nanopore Technology) to gain a more complete picture of the pathogens involved. Throughout the metagenome assembly of a sample harvested from the ear canal of an 11-year-old female Yorkshire terrier suffering from chronic external otitis, a genome of *Lawsonella clevelandensis* was compiled. To our knowledge, this result is the first of its type of animal origin. The outcome of the assembly (CP140010) is a single circular chromosome with a length of 1,878,509 bp, and 1,826 predicted protein-coding genes. No open reading frames associated with antimicrobial resistance could have been identified.

## Introduction

External otitis is a common condition in dogs. The conventional treatment is occasionally rather challenging and often fails. This is partly due to the anatomical characteristics of the ear and partly due to the difficulty in identifying the microorganisms (bacteria, fungi) involved in the development of the disease. Additionally, there is a clear link between the widely occurring food allergy and the prevalence of otitis externa in dogs.^1^ However, other sources of dermal hypersensitivity may also play a role in the clinical scenario. In the case of purely infective external otitis, the lack of information about the drug susceptibility of the pathogens involved in the disease is an additional therapeutic challenge. The above-mentioned factors can lead to so-called end-stage chronic otitis requiring an invasive salvage procedure, the Total Ear Canal Ablation (TECA) with or without Lateral Bulla Osteotomy (LBO).^2^

Classic microbiological tests (e.g., culturing, PCR) can be used as a model-based approach to identify the microorganisms located both in the external ear canal and the tympanic bulla. However, participating bacteria have different growth properties and require different culture conditions or cannot even be cultured using classic techniques, and/or the accompanying microflora may mask the real causative agent or other microbial factors in the multi-microorganism-origin cases. At the same time, next-generation and third-generation sequencing allows a data-driven approach rather than a model-based approach, opening a wider view of the microbiota studied. Digital data generated by partial or complete sequencing of nucleic acids (DNA and/or RNA) extracted from a sample can be compared with known genomes to build up a more exact and real-time cross-sectional microbiological image. Clinical metagenomics^3^, genomic sequencing of microorganisms provides information on their genetic characteristics (e.g., virulence and antimicrobial genes) in addition to their presence and abundance. Moreover, for metagenomic studies, Oxford Nanopore Technology (ONT) takes less time than the conventional microbiological methods.^4^ The data presented here are from a larger series of clinical metagenomic samples from canine otitis externa cases. The sample set is collected to investigate the pathogens involved in the condition. From one of the external auditory canal samples, an essentially complete genome of the emerging pathogenic bacteria *Lawsonella clevelandensis*^5^ could have been generated by metagenome assembly.

## Materials and Methods

### Sample collection

The metagenomic sample was collected from the right external ear canal of an 11-year-old intact female Yorkshire terrier before TECA/LBO surgery. The dog was previously diagnosed with ear canal stenosis due to end-stage external otitis. The duration of signs was more than a year. No tumour tissues were identified via histopathology examination. A presumptive diagnosis was chicken meat allergy, and multiple conservative treatment efforts had been performed previously. The skin of the ear canal was red, swollen, scaled, thickened, and covered with purulent ear discharge before the surgery.

### DNA extraction, library preparation and sequencing

DNA extraction was performed with QIAamp PowerFecal Pro DNA Kit from Qiagen according to the manufacturer’s instructions. The concentrations of the extracted DNA solutions were evaluated with an Invitrogen Qubit 4 Fluorometer using the Qubit dsDNA HS (High Sensitivity) Assay Kit. The metagenomic long-read library was prepared by the Rapid Barcoding Kit 24 V14 (SQK-RBK114.24) from Oxford Nanopore Technologies (ONT). The sequencing was implemented with a MinION Mk1C sequencer using an R10.4.1 flow cell from ONT.

### Bioinformatic analysis

The basecalling was performed using dorado (https://github.com/nanoporetech/dorado, v0.4.3) with model dna_r10.4.1_e8.2_400bps_fast@v4.2.0, based on the POD5 files converted from the FAST5 files generated by the sequencer. The raw reads were adapter trimmed and quality-based filtered by Porechop (v0.2.4, https://github.com/rrwick/Porechop) and Nanofilt (v2.6.0)^6^, respectively. The cleaned reads were taxonomically classified by Kraken2^7^ on a database built (27/11/2023) from the representative bacterial reference genomes obtained from NCBI RefSeq. The reads that were classified to *L. clevelandensis* were subjected to antimicrobial resistance gene (ARG) analysis. All possible open reading frames (ORFs) were predicted by Prodigal (v2.6.3)^8^ on each long-read. The protein-translated ORFs were searched for ARG sequences against the Comprehensive Antibiotic Resistance Database (CARD, v.3.2.8)^9,10^ by the Resistance Gene Identifier (RGI, v6.0.3) De novo assembly was carried out using Flye^11^ (v2.9-b1779), polishing of the contigs was performed with medaka (https://github.com/nanoporetech/medaka, v1.7.2), and scaffolds were created by RagTag (v2.1.0)^12^ using the *L. clevelandensis* representative reference genome (ASM129312v1). The average nucleotide identity (ANI) of scaffolds compared to the genome was estimated by pyani (v0.2.12).^13^ For the genome annotation PGAP^14^ (v2023-10-03.build7061) was used guided, setting the target organism as *L. clevelandensis*. Phylogenetic analysis was performed based on the amino acid sequences of the *transaldolase* gene^15^ using 10 annotated of the 12 available *L. clevelandensis* assembled genomes in the NCBI Genome database (accessed on 16/12/2023). The gene-tree was constructed^16^ based on multiple sequence alignment by MAFFT (v7.490).^17^ The best substitution model was selected by functions of phangorn (v2.11.1) package^18^ based on the Bayesian information criterion. The generated neighbor-joining tree was optimized by the maximum likelihood method. Bootstrap values were produced by 100 iterations. All data processing and plotting were done in the R-environment.^19^

## Results

By taxon classification, 5.67% of the reads (n=5388, length ranged between 116 and 28471 bp, with a median of 1965 bp) were found to be the most similar to *L. clevelandensis* among the representative bacterial reference sequences. The ARGs identified with 100% sequence identity are: *AAC(6’)-Ib8* (coverage: 32.00%), *AAC(3)-Ia* (27.27%), *catA8* (24.65%), *lnuB* (24.34%), *lnuF* (23.08%), *Erm(O)-lrm* (21.54%), *QnrVC7* (17.43%), *baeS* (16.92%), *YojI* (15.17%), *tap* (14.67%), *carA* (13.79%), *poxtA* (13.10%), *tet(42)* (12.62%), *tetB(P)* (12.42%), *ErmX* (10.21%), *OXA-152* (9.82%), *srmB* (9.64%), *lmrP* (6.16%), *Acinetobacter baumannii AbaF* (5.13%), *acrB* (4.77%).

The metagenome-assembled genome (NCBI Accession Number: CP140010) consists of a single circular chromosome with a length of 1,878,509 bp. The comparison of the genome to the reference genome ASM129312v1 resulted in an ANI of 89.36The G+C content of the genome amounted to 58.7%. Overall, 3 complete rRNA and 46 tRNA genes were predicted to be present genome-wide that encode 1,826 proteins. Figure 1 shows the gene-tree based on the amino acid sequences of the *transaldolase* gene with the best substitution model, WAG.

**Figure 1.**
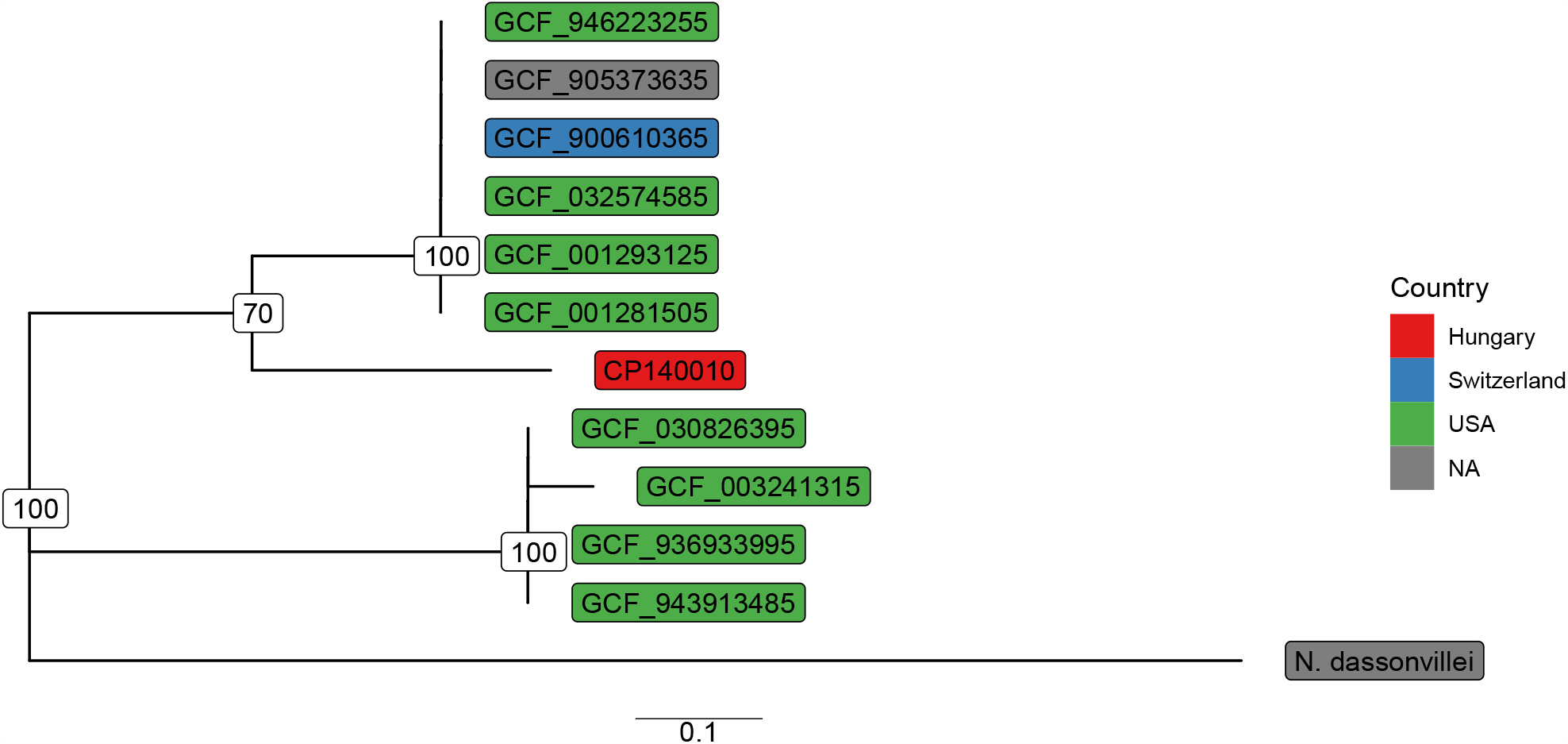
Gene-tree based on *transaldolase* amino acid sequences. RefSeq assembly IDs are shown, and the *Nocardiopsis dassonvillei* (NZ_LR134501.1) was used as an outgroup. Numbers at branches indicate bootstrap support levels (100 replicates). The assemblies originate from abscesses (GCF_001281505, GCF_001293125, GCF_900610365), hospital NICU environment (GCF_003241315), human gut (GCF_032574585), oral (GCF_905373635), and skin metagenomes (GCF_030826395, GCF_936933995, GCF_943913485, GCF_946223255).

## Discussion

*L. clevelandensis* has been detected in multiple human sources associated with health issues, including dermatitis^20^, various abscesses^21–25^, gut dysbiosis^26^, or vascular graft infection^27–29^. Other studies showed that the bacterium can be found in the microbiome of skin^30^, nostrils^31^, human hair follicles^32^, public transit air^33^ or even worn spectacles^34^. Further investigations on the human skin microbiota have revealed interesting findings related to *Lawsonella spp*.. Within a next-generation sequencing-based study, negative correlations among *L. clevelandensis* and the pathological state of human atopic dermatitis were found.^35^

Throughout the work of another research group, *Lawsonella* quantities were found to have a positive correlation with transdermal water loss and a negative correlation with the skin water content. This finding indicates the emergence of this bacterium when the skin integrity is impaired.^36^

To our knowledge, just a few studies have so far shown the presence of *L. clevelandensis* in animals. Within a study, the genome fragments of a member of the *Lawsonella* genus were identified in bull semen by 16S rRNA sequencing^37^. In a further study, *L. clevelandensis* was detected in the oral microbiome of primates using shotgun metagenomics^38^. Furthermore, in a 16S rRNA sequencing study assessing the mucosal microbiome in equine glandular gastric disease, *Lawsonella* genus was found to be associated with the healthy state of the microbiome rather than the pathological conditions.^39^

The NCBI Genome database (accessed on 16/12/2023) has 12 *L. clevelandensis* genomes, all originating from human or human environmental sources. Conclusively, to the best of our knowledge, the presented assembled genome is the first one of animal sources. As demonstrated in Figure 1, sequence identity among *L. clevelandensis* presented in this study, and other strains of the bacterium cannot be explained by sample origins or geographic locations. Importantly, despite the bacterial genome assembly being performed on a sample deriving from an inflammatory environment, the role of the bacterium in external otitis cannot be confirmed. Further studies are required to elucidate its exact contribution to the development and maintenance of the disease. However, a possible role in the pathogenesis of external otitis caused by allergodermatitis may partially be explained by its association with the impairment of the skin barrier function. Further studies are warranted to reveal whether *L. clevelandensis* is a transient microbiota component in dogs, or it is rather of human origin. However, if *L. clevelandensis* is considered as the inhabitant of the dog epidermis, its associations with similar skin changes as in the case of humans would be required to be studied. Importantly, in a recent study,^40^ where the normal microbiota of the external ear canal and middle ear of healthy dogs was examined using 16S rRNA sequencing, *L. clevelandensis* was not found among the constituents of the ear microbiota.

It should also be kept in mind that this often overlooked, opportunistic pathogenic bacterium, which has only recently been described^5^, was identified in the ear canal of a companion animal. At the same time, a change in the quality of human-pet bonds is notable. Current mindset often enhances physical proximity (e.g., sleeping with the owner, unhygienic interactions causing human exposure to canine saliva and other secretions, etc.) among dogs and their owners. Namely, around 70% of owners regard and treat their dogs as family members, 17% as companions, and only 3% as property, according to a survey by the American Veterinary Medical Association.^41^ Considering the nature of modern human-pet bonds^42^, the possibility is raised that this emerging pathogenic bacterium may have zoonotic significance and, thus, enhanced One Health implications.

However, in line with previous studies^43^ that presented no antimicrobial resistance (AMR) determinants identified in the genome of *L. clevelandensis*, along with low to very low Minimal Inhibitory Concentration (MIC) antimicrobial susceptibility results, we have no identified ARG hits with sufficient length and base sequence identity.

*L. clevelandensis* is a fastidious, anaerobic bacterium with a lengthy incubation time of at least 10 days.^22^ As such, conventional culture techniques often fail to isolate the bacterium.^28,29^ Once successfully cultured, the differentiation of *L. clevelandensis* from non-tuberculous mycobacteria is still demanding due to its partial acid resistance.^21^ Therefore, even if diagnostic microbiology laboratories target the pathogen, culturing, and identification during the routine bacteriological testing period is unlikely. Nanopore sequencing and consecutive bioinformatic data processing enable the detection of the microorganisms, combined with the assessment of their abundance rates, virulence genes, and antimicrobial resistance determinants within 4-24 hours.^44^ Thus, ONT-based clinical metagenomics may become an important tool to support diagnostic microbiology and clinical work. Nevertheless, the novelty of this approach leaves many questions unanswered, such as discrepancies between the detected antimicrobial resistance genes and the phenotypic antimicrobial resistance expected.^45^ Similarly, the pathogenic role of microorganisms identified solely by metagenomics in diseases previously associated with one or a few different pathogens necessitates to be comprehended.^46^ Relatedly, pathogenetic correlations of the abundance of detected microorganisms may also open new perspectives in learning, better treating, and hindering the development of diseases with an infectious background in the future.

## Declarations

### Ethics approval and consent to participate

Not applicable.

### Consent for publication

Not applicable.

### Data Availability

The raw long-read data (SRR26949354) of the sample (SAMN38440151) are publicly available and accessible through the PRJNA1045271 from the NCBI Sequence Read Archive (SRA).

### Author contributions statement

NS takes responsibility for the integrity of the data and the accuracy of the data analysis. AGT, NS and TN conceived the concept of the study. TN and ZK collected the biological samples. AGT and NS participated in the bioinformatic analysis. AGT and NS participated in the drafting of the manuscript. AGT, MT, NS, and TN carried out the critical revision of the manuscript for important intellectual content. All authors read and approved the final manuscript.

## Acknowledgements

We thank Sára Ágnes Nagy and Márton Papp for supporting our work.

## Authors’ information

Not provided.

